# Discourse Focus and Memory Encoding: The Role of Trial-Level Alpha Power

**DOI:** 10.1101/2025.07.05.663298

**Authors:** Eleonora J. Beier, Assaf Breska, Lee M. Miller, Yulia Oganian, George R. Mangun, Tamara Y. Swaab

## Abstract

The rapid, continuous flow of spoken language places strong demands on attention, and it is thought that listeners meet these demands by predicting when important information will occur and allocating attention accordingly. However, to date there is little direct evidence for the involvement of preparatory attention during language processing. In this study, we investigate preparatory attention during spoken language comprehension by measuring alpha neural activity with EEG, a known measure of temporal attentional preparation. Alpha activity leading up to target words that were either focused or defocused by a preceding discourse question did not vary as a function of focus, challenging the assumption that attention is pre-allocated to the timing of focused words. On the other hand, we found that trial-by-trial fluctuations in alpha activity predicted both the depth of processing and the subsequent memory for new information. Specifically, pre-target alpha modulated a centro-parietal Dm subsequent memory effect for focused words, linking preparatory attention to memory encoding during comprehension. Together, these findings bridge psycholinguistic studies on information structure and cognitive neuroscience research on temporal attention, offering novel insights into the role of alpha activity in attentional dynamics during spoken language processing.

## Introduction

Spoken language rapidly unfolds over time. Most utterances contain information that is new to the listener, as well as redundant information, organized into what is referred to as the *information structure* of the sentence. Efficient comprehension depends on the ability to identify and integrate new information as it becomes available, making information structure a key organizing principle in real-time processing. The timing of new information can be inferred from the discourse context; for example, given the question “Who brought the cake?”, listeners can predict that the answer to this question will occur early in the following sentence – based on the typical subject-verb-object (SVO) structure of English: “<u>Emma</u> brought the cake”. On the other hand, given the question “What did Emma bring?”, listeners can predict the answer will occur in the later part of the sentence – in the object position. Even in absence of an explicit question, speakers help listeners prepare for upcoming novel information by focusing it through prosody and syntactic focus constructions (e.g., “What she brought was…”). Across languages, listeners are thought to pre-allocate attention to the key points in time when the most important, new information is expected to occur based on these cues (Akker & Cutler, 2003; Beier & Ferreira, 2022; Cutler & Fodor, 1979; Kristensen et al., 2013a; Wang et al., 2014; Yang et al., 2019). Yet despite this theoretical emphasis on attention as a core mechanism, there is surprisingly little direct evidence about how attentional resources are allocated in time during comprehension, or how such mechanisms influence spoken language processing and memory for the discourse.

Non-linguistic studies of temporal prediction have shown that cues in the environment can guide attention to the timing of critical events, a process reflected through modulations in the power of EEG alpha neural activity (8 – 13Hz) (Breska & Deouell, 2017b, 2017a; Nobre & Van Ede, 2018). Alpha activity has also been linked to attention for spoken language, making it a valuable tool for measuring attentional dynamics (Boudewyn et al., 2015; Boudewyn & Carter, 2018; Dimitrijevic et al., 2017; Wöstmann et al., 2016, 2017). Here, we test whether discourse questions – as in the example above – engage temporal prediction mechanisms to direct attention towards focused words. Specifically, we examine preparatory attention by measuring alpha power *prior* to the onset of words that were focused by the discourse, and test whether such preparatory states influence how these words are processed and remembered.

### Information structure and attention

Many studies show that prediction is a fundamental feature of language processing and cognition more broadly (Clark, 2013; Federmeier, 2007; Ferreira & Chantavarin, 2018; Kuperberg & Jaeger, 2016; Pickering & Garrod, 2007). Most of this work has focused on continuous predictions about the *content* of upcoming information, such as semantic and syntactic features (Kuperberg & Jaeger, 2016; Pickering & Garrod, 2007). Less attention has been devoted to how comprehenders predict the *timing* of that information. While temporal predictions are known to facilitate perceptual and motor processing in non-linguistic domains (Nobre & Van Ede, 2018), their role in guiding attention during language comprehension remains poorly understood (Kotz & Schwartze, 2010).

Psycholinguistic studies show that cues to the timing of upcoming important information have profound effects on comprehension across modalities. Early studies found that listeners were faster at detecting phonemes at the start of words focused either through the preceding discourse question (Beier & Ferreira, 2022; Cutler & Fodor, 1979) or the speaker’s prosodic intonation (Akker & Cutler, 2003; Cutler, 1976). Subsequent ERP studies found that words focused by a preceding question and/or through prosodic pitch accents elicited a larger N400 to semantic violations (Wang et al., 2009, 2011) and a larger P600 to syntactic violations (Wang et al., 2012), reflecting increased sensitivity to meaning and grammatical structure, suggesting deeper linguistic processing. This is in line with eye-tracking reading studies, showing longer reading times for words focused through the discourse and through syntactic focus constructions (S. Birch & Rayner, 1997; Lowder & Gordon, 2015). This increased depth of processing is likely responsible for the robust effect of focus on memory: comprehenders were consistently more likely to remember words when they had been focused through the discourse (Cutler & Fodor, 1979), prosody (Fraundorf et al., 2010, 2012) or syntax (S. L. Birch & Garnsey, 1995).

While this evidence establishes that information structure plays a crucial role across all layers of language processing – from phoneme detection, to semantic and syntactic processing, to later memory – the underlying cognitive and neural mechanisms remain debated. A prevalent explanation for these effects is that listeners anticipate the timing of focused words and pre-allocate attention and processing resources to those points in time, enhancing their ability to process the information efficiently (Akker & Cutler, 2003; Beier & Ferreira, 2022; Cutler & Fodor, 1979; Wang et al., 2014). Although this explanation is widely invoked, most studies have not provided direct evidence for the underlying attentional mechanisms. However, there is some limited evidence suggesting that attention plays a role. Using fMRI, Kristensen et al. (2013) found that words focused through a pitch accent involved activation of the bilateral superior/inferior parietal cortex, superior temporal cortex, and left precentral cortex, a network of regions associated with domain-general attention (Corbetta et al., 2000, 2008; Giesbrecht et al., 2003; Hopfinger et al., 2000; for a review, see Mangun et al., 2025). Similarly, words focused through the discourse (Yang et al., 2019) and through syntax (Chen et al., 2014) led to a larger P2, an ERP response associated with attention (Luck & Hillyard, 1994; Singhal et al., 2002).

However, these studies only observed attentional effects *following* the presentation of the focused word. To the best of our knowledge, there is currently no direct evidence for the hypothesized pre-allocation of attention *prior* to a focused word, resulting from a temporal prediction.

Furthermore, there are alternative explanations for the behavioral and ERP effects of information structure on comprehension that do not necessarily involve attention. It has been suggested that information structure influences semantic prediction, such that comprehenders are more likely to semantically predict upcoming focused words while engaging in good-enough processing for defocused (less relevant) information (Ferreira & Lowder, 2016). This could explain prior findings on the N400, as greater semantic prediction would result in a larger N400 to semantic violations (as in Wang et al., 2011). Greater prediction of focused words could also result in faster phoneme detection (i.e. if phonological information was partially pre-activated; Kuperberg & Jaeger, 2016) and memory effects (Hodapp & Rabovsky, 2021; Hubbard et al., 2019). Thus, prior effects of focus on language processing could arise entirely through modulations in the strength and content of semantic prediction, without requiring the involvement of attentional processes.

Overall, effects of information structure on comprehension have often been explained in terms of attentional pre-allocation to upcoming words, guided by cues from the preceding discourse, prosody, and syntactic structure, that signal when novel information is likely to occur. However, there is little direct evidence for the involvement of attention, and it has not yet been established whether attention is pre-allocated in a predictive manner. To examine this, we use insights from non-linguistic studies of temporal attention, which reveal that preparatory attention to the timing of an upcoming stimulus can be measured through modulations in the power of neural activity in the alpha band prior to the stimulus (Breska & Deouell, 2017b, 2017a; Nobre & Van Ede, 2018).

While alpha and temporal attention have been primarily studied using non-linguistic stimuli (across both the visual and auditory modalities), alpha power has also been shown to reflect auditory attention to speech. Studies that manipulated attention on a global level – attending or suppressing an entire speech stream – found greater alpha power over parietal and temporal regions as listeners suppressed a distractor sentence (Wöstmann et al., 2017); this effect was lateralized to the hemisphere ipsilateral to the attended stream during dichotic listening (Wöstmann et al., 2016). Attention to speech may involve both the inhibition of task-irrelevant regions through alpha synchronization (leading to greater power), as well as a – less robust – pattern of alpha desynchronization (lower power) reflecting greater neural activation of relevant regions (Dimitrijevic et al., 2017), following the gating-by-inhibition view of alpha (Jensen & Mazaheri, 2010). There is also evidence that alpha power reflects local fluctuations in attention towards a single speech stream. An increase in alpha at central sites was shown during attentional lapses (mind-wandering episodes) in naturalistic listening (Boudewyn & Carter, 2018). Similar effects were found in psycholinguistic paradigms, where trial-level alpha fluctuations predicted ERP responses to referential ambiguity (Nref; Boudewyn et al., 2015) and semantic incongruity (N400; Boudewyn et al., 2017). Alpha was also shown to predict behavioral performance and memory for linguistic content (Boudewyn & Carter, 2018; Dimitrijevic et al., 2017).

Taken together, these studies suggest that alpha power is a reliable indicator of attention during spoken language processing, linked to ERP and behavioral measures of comprehension. In this study, we employ this measure to examine the claims of prior psycholinguistic studies on information structure, testing whether attention is pre-allocated to the timing of words focused by the prior discourse context, and how this affects comprehension and memory.

### The present study

This study bridges the psycholinguistic literature on information structure, non-linguistic studies of temporal attention, and the body of work examining alpha power as a neural marker of attention during speech processing. We directly test the predominant hypothesis that listeners pre-allocate attention to upcoming focused words, by tracking attention allocation over the course of the sentence through alpha power dynamics.

We employed a discourse question-answer paradigm, adapted from prior behavioral work showing faster phoneme detection for focused words (previously interpreted in view of attention; Beier & Ferreira, 2022; Cutler & Fodor, 1979). On each trial, participants heard either an Early focus question (“Which man was wearing the hat?”) or a Late focus question (“What hat was the man wearing?”), focusing either an Early target or a Late target word in the following sentence (“The man on the <u>corner</u> was wearing the <u>dark</u> hat”; target words are underlined), while EEG was continuously recorded (see Fig. 1). A subsequent memory test measured participants’ memory of the target words (either the Early or Late target for each trial), as a function of whether they had been focused or defocused by the preceding question. We expected to replicate prior behavioral findings of greater memory for focused words (S. L. Birch & Garnsey, 1995; Cutler & Fodor, 1979; Fraundorf et al., 2010).

**Figure 1.**
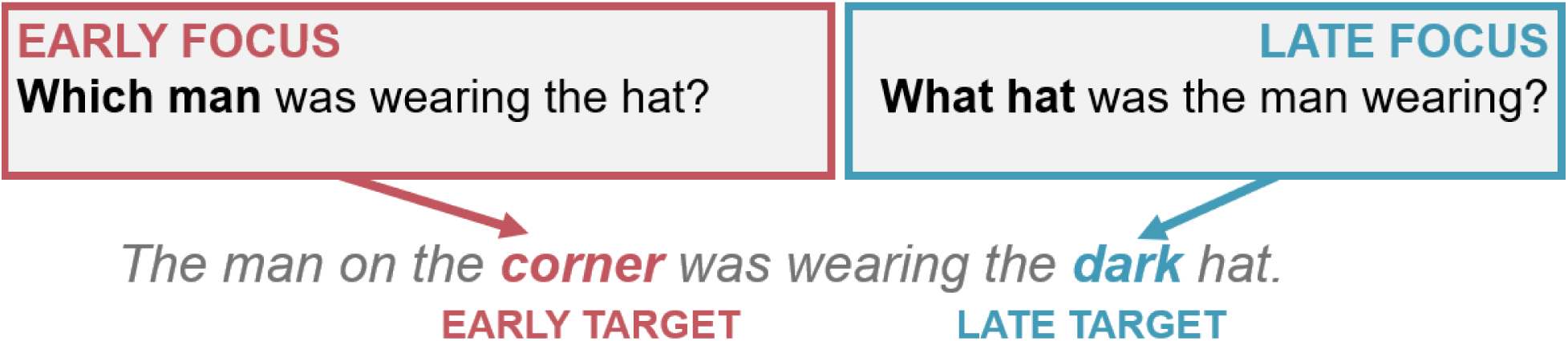
Example stimulus, showing the experimental design. The Early target word is focused when preceded by the Early focus question and defocused when preceded by the Late focus question, and vice versa for the Late target word.

Based on previous studies of attentional fluctuations within a single speech stream (Boudewyn et al., 2015, 2017; Boudewyn & Carter, 2018), we expected to find lower alpha power at central sites prior to the onset of focused, relative to defocused target words, reflecting attention pre-allocation. If discourse focus does not modulate alpha power prior to focused words, this would speak against the engagement of temporal attentional preparation, and would suggest instead that information structure affects processing through language-specific, non-attentional mechanisms such as increased semantic prediction. To further examine whether semantic processing was affected by information structure, we tested whether there was an effect of focus on the N400 elicited by the target words, as prior work showed a larger N400 effect to semantic violations for focused words (Wang et al., 2012). While the target words in our study were not manipulated for predictability or semantic congruence, prior research has shown that the N400 is elicited by every content word in a sentence, and its amplitude is sensitive to processing demands, task goals and contextual support (Bentin et al., 1993; Van Petten & Kutas, 1990). Thus, we predicted that the amplitude of the N400 would vary as a function of discourse focus, reflecting either facilitated semantic processing (a smaller N400 amplitude) or deeper processing (larger amplitude) for focused words.

Further, we addressed whether pre-target alpha and ERP responses during the presentation of the target words predicted accuracy in the subsequent memory test, in order to explore the functional mechanisms behind prior findings of greater memory for focused words. Based on prior work, we expected to find that lower pre-target alpha at central sites (greater attention pre-allocation) would result in better subsequent memory (Boudewyn & Carter, 2018). This would also match studies relating alpha during encoding to later recall in what is referred to as a “difference due to memory” (Dm) effect (Klimesch et al., 1996). Dm effects also include several distinct ERPs, including frontal and parietal positive components at various latencies, which predict subsequent memory (Friedman & Johnson Jr., 2000; Mecklinger & Kamp, 2023; Paller et al., 1987). These components are thought to broadly reflect memory encoding processes (for a review, see Mecklinger & Kamp, 2023). Thus, we tested for potential ERP Dm effects in our design, addressing whether memory encoding of the target words was affected by the discourse focus and by pre-target preparatory attention.

## Method

### Transparency and openness

The method, analyses and hypotheses of this study were pre-registered (AsPredicted #133353) prior to the start of data collection.

### Participants

A total of 48 participants took part in the experiment. All participants provided informed consent. Data from 8 participants were removed from the analyses following one of our pre-registered exclusion criteria: the participant did not complete the experiment (*N* = 1); technical issues during data collection (*N* = 5); or more than 25% of trials rejected due to artifacts, for any of the primary experimental conditions (*N* = 2). This resulted in our pre-registered sample size of 40 participants (26 female, 12 male, 2 non-binary). The median age was 20 years (*SD* = 3.39).

All participants were native English speakers (defined as having learned English before age 5). Participants reported having no hearing, speech or language impairments and no neurological disorders. Further, all participants were right-handed (Grey et al., 2017). Participants were recruited through a University of California, Davis participant pool and were compensated with either class credit or cash ($15 per hour).

### Stimuli

Stimuli consisted of 80 question-answer pairs, including the 36 stimuli used in (Beier & Ferreira, 2022) and another 44 stimuli modeled after those used by (Cutler & Fodor, 1979).

Each stimulus consisted of an Early focus question (“Which man was wearing the hat?”) and a Late focus question (“What hat was the man wearing?”), focusing either an Early target or a Late target word in the following sentence (“The man on the <u>corner</u> was wearing the <u>dark</u> hat”; target words underlined). As in (Cutler & Fodor, 1979) phoneme monitoring experiment, all target words started with one of three possible phonemes (/b/, /d/ or /k/); while this study did not involve a phoneme monitoring task, we maintained this design feature for secondary analyses related to phonological processing, which are beyond the scope of this paper.

All stimuli were recorded by a male native English speaker from California and digitized at 48000 Hz (later downsampled to 44100 Hz) using a Shure KSM44 microphone. As in Cutler and Fodor (1979) and Beier and Ferreira (2022), the speaker was instructed to read each sentence with natural intonation without placing particular emphasis on either the Early or Late target words, to avoid potential confounds of prosodic focus marking. Stimuli were amplitude normalized at 65 dB using a custom Praat script (Boersma & Weenink, 2022). The onset of all words during the sentences, including the Early and Late target words, was determined through the Montreal Forced Aligner (McAuliffe et al., 2017).

#### Filler Items

Stimuli also included 100 filler items and 18 practice items used during an initial practice block. The filler items served several purposes: (1) 48 filler items (26% of all stimuli) were followed by comprehension questions to encourage participants to listen for comprehension (see *Procedure*); we chose to use filler items rather than the experimental items because the repetition involved in answering these questions could influence later memory for the target words, potentially affecting the results. (2) To prevent participants from noticing that the memory test exclusively focused on the Early/Late target words in the experimental items (which were always subject or object modifiers), the memory probes for the filler items targeted other types of words (e.g., verbs, head nouns, prepositions, etc.) (3) Filler items were further designed to provide some variability in the sentence structure to prevent participants from adopting a shallow-processing strategy due to repeated exposure to similar constructions (Bock, 1986; Pickering & Ferreira, 2008), which has been shown to influence ERP responses (Ledoux et al., 2007; Tooley et al., 2009); (4) Finally, 60 filler items contained a manipulation of semantic constraint. Because our pre-registered hypotheses included a potential effect of discourse focus on the N400, these filler items allowed us to assess whether our experimental setup could reliably detect the well-established effect of semantic prediction on this ERP.

The 60 filler items were adapted from a prior study on the effect of semantic constraint on the N400 (Brothers et al., 2019). There were two versions of each sentence: a High Constraint version leading up to a highly predictable target word (e.g., “The bride refused to invite her father to the <u>wedding</u> for some reason”; target word underlined); and a Low Constraint version where the same target word was less predictable, yet plausible (“The lazy teenager refused to go to the <u>wedding</u> for some reason”). The original study (Brothers et al., 2019) ensured the effectiveness of this manipulation through a cloze norming task, where participants read each sentence frame (e.g., “The bride refused to invite her father to the…”) and write in the first word that comes to mind; cloze probability is the percentage of participants completing the sentence with the target word. This confirmed that the target word was more predictable in the High Constraint (95% cloze probability) than in the Low Constraint condition (>1%). To prevent participants from noticing a difference between these filler items and the experimental items and to maintain a similar trial structure, each sentence was preceded by a question (e.g., “Who refused to do something?”). The questions were designed not to focus the semantic target word (e.g., “wedding”) to avoid confounding the effects of discourse focus and semantic constraint.

Additionally, the questions were constructed to avoid increasing the predictability of the subsequent sentence. The remaining 40 filler items were adapted from Beier and Ferreira (2022) and were all followed by comprehension questions. Importantly, the proportion of Early and Late focus questions across all filler and practice items was 50:50. The filler and practice items were recorded by the same native English speaker as the experimental items and amplitude normalized at 65 Hz.

### Procedure and Apparatus

After providing informed consent, participants completed a demographic survey. Following EEG capping (see *EEG Recording*), participants sat in an electrically shielded, sound-attenuated booth in front of a VIEWPixx/EEG LED monitor (model VPX-2006A; VPixx Technologies Inc.) with a refresh rate of 120 Hz. Auditory stimuli were presented through earphones (ER-1, Etymotic Research, Elk Grove Village, IL, USA) via air-tubes, in order to minimize electrical interference with the EEG. After receiving written and verbal instructions about the task, participants completed a practice block.

Stimuli were presented using Presentation software (Version 23.0, Neurobehavioral Systems, Inc., Berkeley, CA). Participants started each trial by pressing the spacebar. After 1000 ms, they heard one of the focus questions (Early or Late). The ISI between the end of the question and the onset of the sentence was jittered between 1180 and 2200 ms to prevent participants from forming a precise temporal prediction beyond the Early/Late manipulation. Each trial ended 1000 ms after the end of the sentence, after which participants could proceed to the next trial by pressing the spacebar. Participants were instructed to fixate on a central cross throughout each trial and to minimize blinking to reduce eye movement artifacts in the EEG. They were encouraged to use the self-paced breaks between trials to blink and rest their eyes.

On 48 filler trials (26% of all trials), the end of the sentence was followed by a question mark presented at the center of the screen. These trials served as comprehension questions: participants were instructed to repeat aloud the answer to the question they just heard. For example, after hearing the question “Which rabbit was chewing the carrot?” followed by the sentence “The white rabbit was chewing the delicious carrot”, participants should respond with: “the white rabbit“. Participants had 5 seconds to answer the question, and their responses were recorded. This task both encouraged participants to use the Early/Late Focus questions as cues for when to pay particular attention during the sentence and ensured that they remained attentive throughout the experiment.

Stimuli were presented in 10 blocks of 18 trials each; each block lasted approximately 5 minutes. Blocks ended with a memory test similar to that used by Cutler and Fodor (1979) and Beier and Ferreira (2022). We chose to test memory after each block rather than at the end of the entire experiment (as in those previous studies) due to the increased number of stimuli and duration of the study. During the memory test, each sentence was presented visually on the screen with one target word blocked out. Participants were instructed to select the correct word completing the sentence among four alternatives. For the experimental items, half of the judgments were about the Early target and half about the Late target words; half were for words that had been focused and half defocused by the preceding questions. For the filler items, memory was tested for other parts of the sentence (e.g., verbs, head nouns, prepositions) to prevent participants from developing a strategy based on the typical position of the probed words.

The experimental conditions (Early/Late focus question, Early/Late target tested in the memory test) were counterbalanced across four lists. The lists also counterbalanced the High Constraint and Low Constraint conditions for the 60 semantic constraint filler items. Stimulus order was pseudo-randomized within each block so that no more than 3 stimuli in the same Focus condition and no more than 2 stimuli of the same type (experimental or filler) were presented consecutively. In addition, each block always started with two filler items and ended with a filler item, so that the experimental items were never at the very beginning or end of the block. Participants were encouraged to take longer breaks between blocks, to reduce fatigue and maintain focus.

### EEG Recording

EEG was recorded using a 64-channel actiCAP active electrode system (Brain Products GmbH) at the following recording sites following the International 10–10 system (Jurcak et al., 2007): FP1, FP2, AF7, AF3, AFz, AF4, AF8, F7, F5, F3, F1, Fz, F2, F4, F6, F8, FT9, FT7, FC5, FC3, FC1, FCz, FC2, FC4, FC6, FT8, FT10, T7, C5, C3, C1, Cz, C2, C4, C6, T8, TP9, TP7, CP5, CP3, CP1, CPz, CP2, CP4, CP6, TP8, TP10, P7, P5, P3, P1, Pz, P2, P4, P6, P8, PO7, PO3, POz, PO4, PO8, PO9, O1, Oz, O2, PO10. Additionally, bipolar EOG channels were used to capture blinks and horizontal eye movements through passive tin electrodes^1^. Data were amplified through a Neuroscan SynAmps2 input board and amplifier and digitized at 1000 Hz using CURRY 8 acquisition software (Compumedics USA, Inc.). Electrode AFz was used as the ground, and electrode FCz as the reference during recording. Data was later re-referenced to the algebraic average of electrodes TP9 and TP10 (placed over the left and right mastoids, respectively) during data pre-processing. The fitted elastic electrode caps were centered for each participant so that electrode Cz was at the half-way point between nasion and inion, and between preauricular points. Electrode impedances were kept below 15 kΩ through the application of high viscosity electrolyte gel at each electrode site.

### EEG Pre-Processing

Data were pre-processed in MATLAB (Mathworks) using the EEGLAB Toolbox (Delorme & Makeig, 2004) and ERPLAB plugin (Lopez-Calderon & Luck, 2014). Pre-processing scripts were adapted from the standardized pre-processing pipeline recommended by (Boudewyn et al., 2023) and available on the Open Science Framework (https://osf.io/wdrj3/). Data from all 10 blocks were first concatenated for each subject, and noisy or broken channels were marked through visual inspection. Independent Component Analysis (ICA) was used on the continuous data to remove the contribution of blink artifacts to the EEG. Data were filtered between 0.05 Hz and 30 Hz using the EEGLAB function *pop_basicfilter*, and re-referenced to the mastoids. Data from noisy or broken channels were interpolated as the average of neighboring electrodes.

For ERP analyses, EEG data were epoched from −200 ms to 1000 ms relative to the onset of the target words, and baseline correction was applied using the 200 ms prior to word onset. For time-frequency analyses, data were epoched from −3000 ms to 3000 ms relative to the onset of the target words and other events of interest (e.g., question and sentence onsets). For these epochs, we did not apply baseline correction to the EEG voltage at this point, given that we were interested in oscillatory activity before target word onsets (see *Time-Frequency Analyses* for more details on the baseline correction later applied to spectral power). These longer epochs were used to measure oscillatory activity over the periods before and after the target words, and to account for edge artifacts, which occur when wavelet convolution extends beyond the available data at the edges of each epoch, potentially distorting power estimates.

Finally, artifact rejection was performed separately for the shorter and longer epochs using the peak-to-peak algorithm through the ERPLAB function *pop_artmwppth*, starting with an amplitude threshold of 100 µV which was iteratively adjusted for each participant through visual inspection (resulting in a range from 60 to 160 µV).

## ERP Analyses

Following our pre-registered hypotheses, we initially extracted mean amplitude over a window of 300-500 ms following target word onset, matching the typical latency of the N400 (Šoškić et al., 2022). We also performed follow-up analyses on a later time window of 600-1000 ms. Mean amplitude was calculated on a trial-by-trial basis for each participant, and analyzed through mixed effects models using the *lme4* and *lmerTest* packages in R (Bates et al., 2014; Kuznetsova et al., 2017). We used mixed effects models rather than ANOVAs to more flexibly account for by-subject and by-item variability, and because this analytical method has been shown to be more powerful (Baayen et al., 2008; Stevens & Brysbaert, 2016) and more robust to noise, specifically for ERP data (Heise et al., 2022). To account for differences in the scalp distribution of the effects, we included a fixed effect of electrode cluster (Central: FC1, FC2, C1, CP1, CPz, C2, Cz, CP2; Left-Frontal: F7, F3, FC5, FT7, FC3, Fp1, F5, AF7; Left-Posterior: PO7, PO9, P5, P3, O1, P7, PO3; Left-Temporal: TP7, CP3, C3, C5, T7, CP5; Mid-Frontal: Fz, AF4, AFz, F2, AF3, F1; Mid-Posterior: P1, Pz, Oz, POz, P2, Iz; Right-Frontal: F6, F4, FT8, FC4, AF8, F8, FC6, Fp2; Right-Posterior: PO4, P4, PO8, P6, O2, PO10, P8; Right-Temporal: CP4, T8, C6, C4, CP6, TP8). The effect of cluster was dummy-coded so that contrasts were performed against the Central cluster.

The model on the semantic constraint filler items included a fixed effect of semantic constraint (High, Low; dummy-coded) and its interaction with cluster, as well as by-subject and by-item random intercepts and slopes for semantic constraint.

The model on the experimental items was performed separately for the Early and Late target words. Models included a fixed effect of focus (Focused, Defocused; dummy-coded) and its interaction with cluster, as well as by-subject and by-item random intercepts and slopes for focus.

Lastly, we tested whether ERP mean amplitude varied as a function of pre-target alpha on a trial-by-trial basis, aggregating across the Early and Late targets. The model included fixed effects of focus (Focused, Defocused) and pre-target alpha (see *Time-Frequency Analyses*), as well as by-subject and by-item random intercepts and slopes for focus. For this model, the effect of focus was sum-coded in order to estimate the main effect of pre-target alpha across focus conditions.

### Time-Frequency Analyses

Event-related spectral power was estimated using the EEGLAB function *newtimef*, relative to the onset of the Early and Late target words. Power was estimated for a range of linearly spaced frequencies from 3 to 30 Hz using Hanning-tapered wavelets starting at 3 cycles and increasing in cycles for higher frequencies. Baseline power calculated over the 1000 ms prior to the focus question was log-subtracted using a common baseline; we chose this as our baseline window given that any differences induced by the focus condition could in principle begin during the focus question. Alpha power was calculated as the average power between 8 and 13 Hz. Alpha power was compared between the Focused and Defocused conditions in the 1500 ms prior and following each target word, using cluster-based permutation statistics. Our initial analysis was conducted on electrode Cz, where we expected to find lower alpha power corresponding to greater preparatory attention based on Boudewyn et al. (2015, 2017) and Boudewyn and Carter 2018). Follow-up analyses were performed on a range of Central and Occipital midline electrodes (CPz, Pz, POz, Oz)^2^.

We also performed follow-up analyses on trial-level pre-target alpha, estimated using the BOSC method (Whitten et al., 2011) through the fBOSC toolbox in MATLAB (Seymour et al., 2022). Similarly to Boudewyn and Carter (2018), we employed this approach to address an important limitation: given the variable duration and number of words across stimuli, there is likely to be jitter in the exact onset of any differences in alpha power across conditions. Thus, average alpha power averaged across trials may obscure an effect that has inconsistent timing across stimuli. The BOSC method overcomes this issue by calculating the percentage of timepoints over a window (1500 ms prior to target word onset) that contain periodic activity within the alpha band above and beyond background 1/f aperiodic activity. In the fBOSC toolbox, 1/f aperiodic activity is calculated using the FOOOF algorithm (Donoghue et al., 2020). We then detected episodes of periodic activity exceeding a power threshold (above the 95th percentile of the theoretical probability distribution) and duration threshold (3 cycles per frequency), for each frequency, averaging over the alpha band (8 - 13 Hz). The resulting *p-episode* metric therefore represents the percentage of timepoints over the selected window that contain alpha activity above these thresholds, regardless of its exact timing across each stimulus. We used this measure of pre-target alpha as a trial-by-trial predictor when analyzing ERP mean amplitudes and behavioral measures.

We also tested whether trial-level pre-target alpha varied as a function of focus condition. To deal with the non-normal, zero-and-one-inflated distribution of the p-episode metric, we used a Bayesian ordered-beta regression through the *ordbetareg* package in R (Kubinec, 2023). Similarly to the ERP analyses, the model was conducted separately for the Early and Late target and included fixed effects of focus (Focused, Defocused; dummy-coded), cluster, and their interaction. It also included by-subject and by-item random intercepts and slopes for focus.

### Behavioral analyses

Accuracy on the memory test was analyzed through a logistic mixed effects model with fixed effects of Target Position (Early, Late), Focus Position (Early, Late) and their interaction. The model included by-subject and by-item intercepts and slopes for Target Position and Focus Position.

We also performed an exploratory analysis on the time it took participants to make their selection on the memory test. This included the time it took participants to read the sentence (with the target word blocked out) and the four possible options, and participants were not instructed to be as fast as possible. Based on the response time distribution, response times faster than 2 seconds and slower than 30 seconds were excluded from the analysis. Given that the distribution was right-skewed, response times were also log-transformed to achieve a normal distribution (Baayen & Milin, 2010). Response times were then analyzed through a linear mixed effects model with the same fixed and random effects as the model on accuracy. As the full model did not converge, we removed the random slope of Focus Position^3^.

We additionally tested whether memory test accuracy was predicted by pre-target alpha and ERP mean amplitude over the 600-1000 ms window, on a trial-by-trial basis. Data was aggregated across the Early and Late target words. We used a logistic mixed effects model with fixed effects model with fixed effects of focus (Focused, Defocused), pre-target alpha and ERP mean amplitude. The effect of focus was sum-coded, while pre-target alpha and ERP mean amplitude were z-scored to help with model convergence. The model also included by-subject and by-item random intercepts.

## Results

### Behavioral results

On average, participants performed with 77% accuracy (*SD* = 42%) on the memory test (similarly to prior studies; Cutler & Fodor, 1979). A logistic mixed effects model found that accuracy was overall higher for the Early than the Late target word (β = −0.81, *SE* = 0.21, *p* < 0.0001), and overall higher for the Early than the Late focus condition (β = −0.86, *SE* = 0.15, *p* < 0.0001). Critically, there was a significant interaction, indicating that memory for each target word was higher when it had been focused by the preceding question (β = 1.61, *SE* = 0.19, *p* < 0.0001; Fig. 2), replicating Cutler and Fodor (1979).

**Figure 2.**
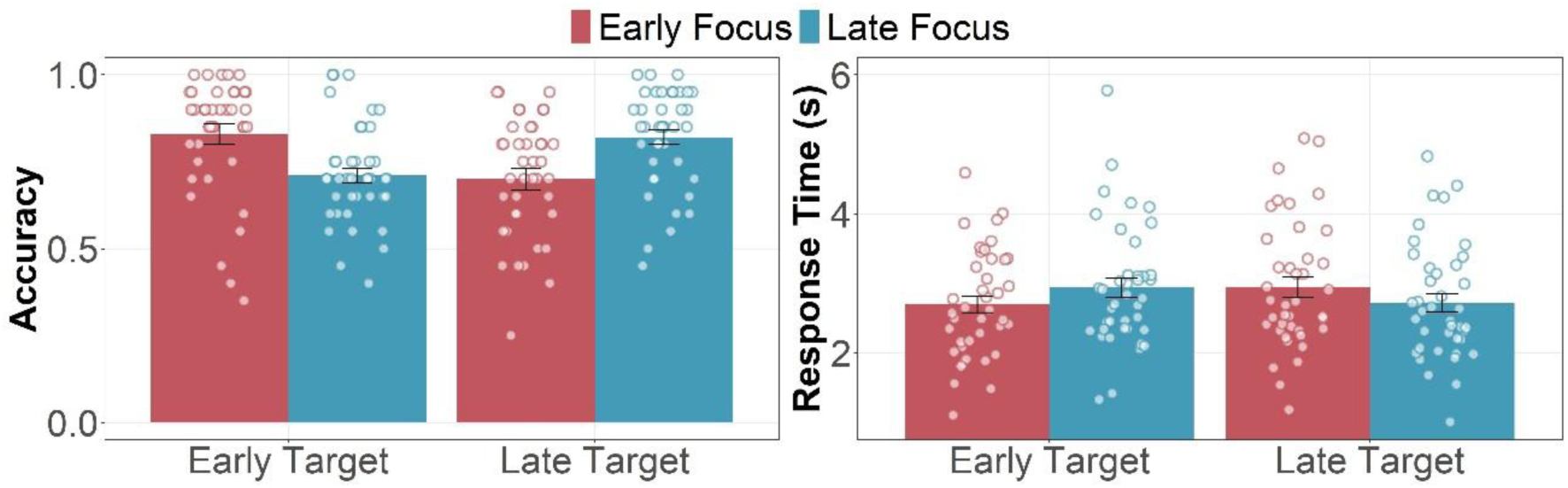
Memory test accuracy and response times, for Early versus Late target words, as a function of Early versus Late focus. Greater accuracy and faster response times were found for the Early target when focused via the Early focus question, and vice versa for the Late target. Data points represent mean accuracy and response times for each participant. Error bars represent the Standard Error of the Mean.

The exploratory analysis on response times in the memory test revealed that participants were overall faster at selecting an answer for the Early than the Late target word (β = 0.1, *SE* = 0.02, *p* < 0.0001), and overall faster for the Early than the Late focus condition (β = 0.09, *SE* = 0.02, *p* < 0.0001). Like for accuracy, there was a significant interaction, indicating that participants were faster for words that had been focused by the preceding question (β = −0.17, *SE* = 0.3, *p* < 0.0001).

Thus, we found that participants were both more accurate and faster at selecting a response to the memory test for words that had been focused during the trials. We interpret the faster selection as reflecting a greater amount of confidence in their response.

### ERP results

#### Effect of semantic constraint

As expected, central electrodes showed the N400 effect, that is a stronger negative deflection for Low Constraint as compared to High Constraint filler items 400 ms past word onset (effect of constraint at 300-500 ms: β = −2.26, *SE* = 0.4, *p* < 0.0001, Fig. 3). Significant interactions with cluster confirmed that the difference between High and Low Constraint was significantly greater at Central sites compared to other electrode clusters (see Table S1 for full model output). This analysis establishes that our design had sufficient power to detect the effect of semantic constraint on the N400.

**Figure 3.**
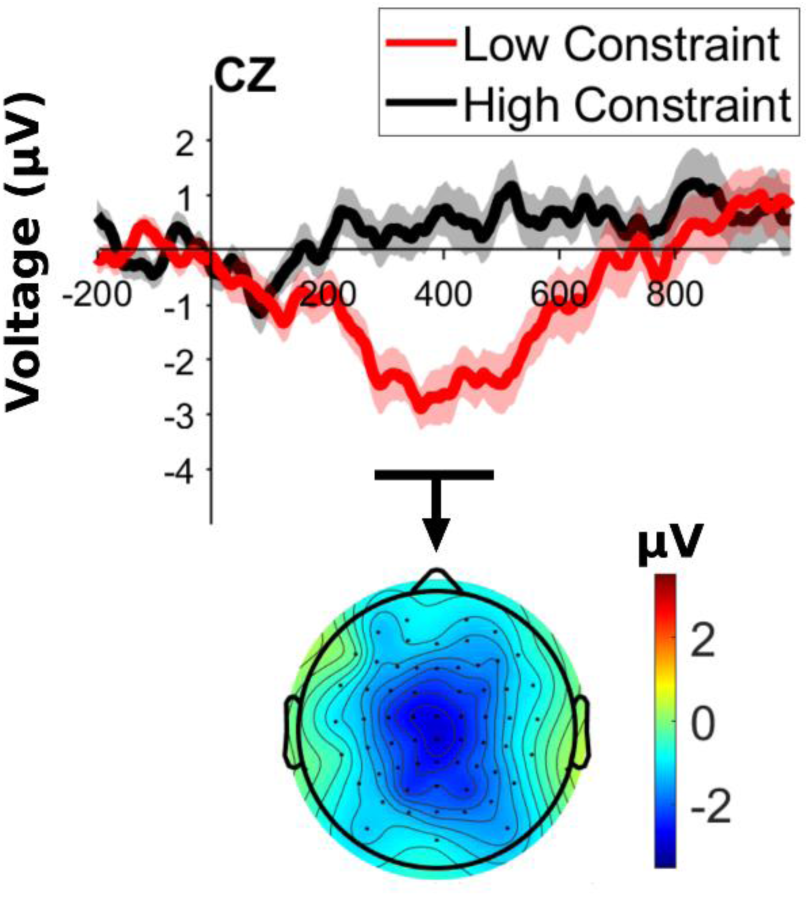
ERP at electrode Cz, locked to the onset of the target word in the semantic constraint filler items, as a function semantic constraint. The scalp distribution represents Low minus High Constraint, averaged over 300-500 ms post-target. Note that positive is plotted upwards on the y-axis for all ERP plots.

#### Effect of focus

For the experimental items, we did not observe a modulation in the N400 as a function of focus (no significant effect of focus for either Early or Late targets at 300-500 ms, *p* > 0.05; Fig. 4; see Table S2 for full model outputs). Instead, for both the Early and the Late target words we observed a late positivity for Focused as compared to Defocused target words. The effect of focus at 600-1000 ms was significant for the Late target (β = 1.08, *SE* = 0.33, *p* < 0.001) and marginally significant for the Early target (β = 0.69, *SE* = 0.39, *p* = 0.08). The effect was maximal at Central and Posterior sites (see Table S3 for full model output).

**Figure 4.**
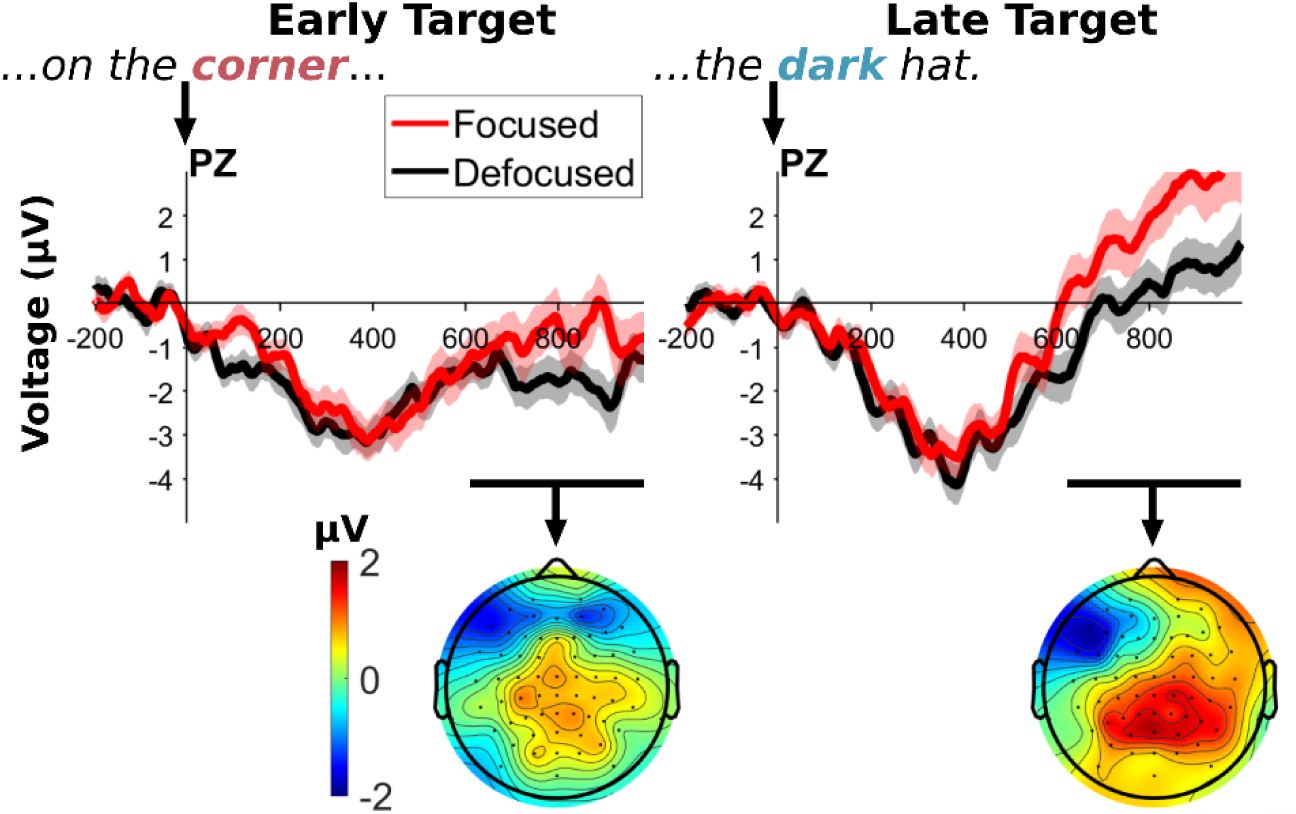
ERPs at electrode Pz, locked to the onsets of the Early and Late target words, as a function of focus condition. The scalp distribution represents Focused minus Defocused conditions, averaged over 600-1000 ms post-target.

In sum, we did not see an effect of focus on the N400; rather, we found that focused words elicited a centro-parietally distributed late positivity relative to defocused words. Late positivities, particularly those with a centro-parietal distribution following the N400, are often associated with stimulus evaluation, semantic integration, and depth of encoding. Such components have been linked to strategic processing and enhanced memory for task-relevant information (Mecklinger & Kamp, 2023). Given that focused words elicited a robust late positivity in our study, we sought to determine whether this effect reflected deeper encoding processes— indexed by improved memory performance—and whether it was modulated by preparatory brain states, as indexed by pre-target alpha power.

#### Alpha results

Event-related spectral power in the alpha band is shown in Fig. 5. Contrary to our predictions based on prior studies (Boudewyn et al., 2015, 2017; Boudewyn & Carter, 2018), the cluster-based permutation did not reveal lower pre-target alpha power in the Focused condition at electrode Cz for either target. We did, however, find a significant cluster at Cz between 960 - 1303 ms relative to the Late target word onset, with lower alpha in the Focused than the Defocused condition.

**Figure 5.**
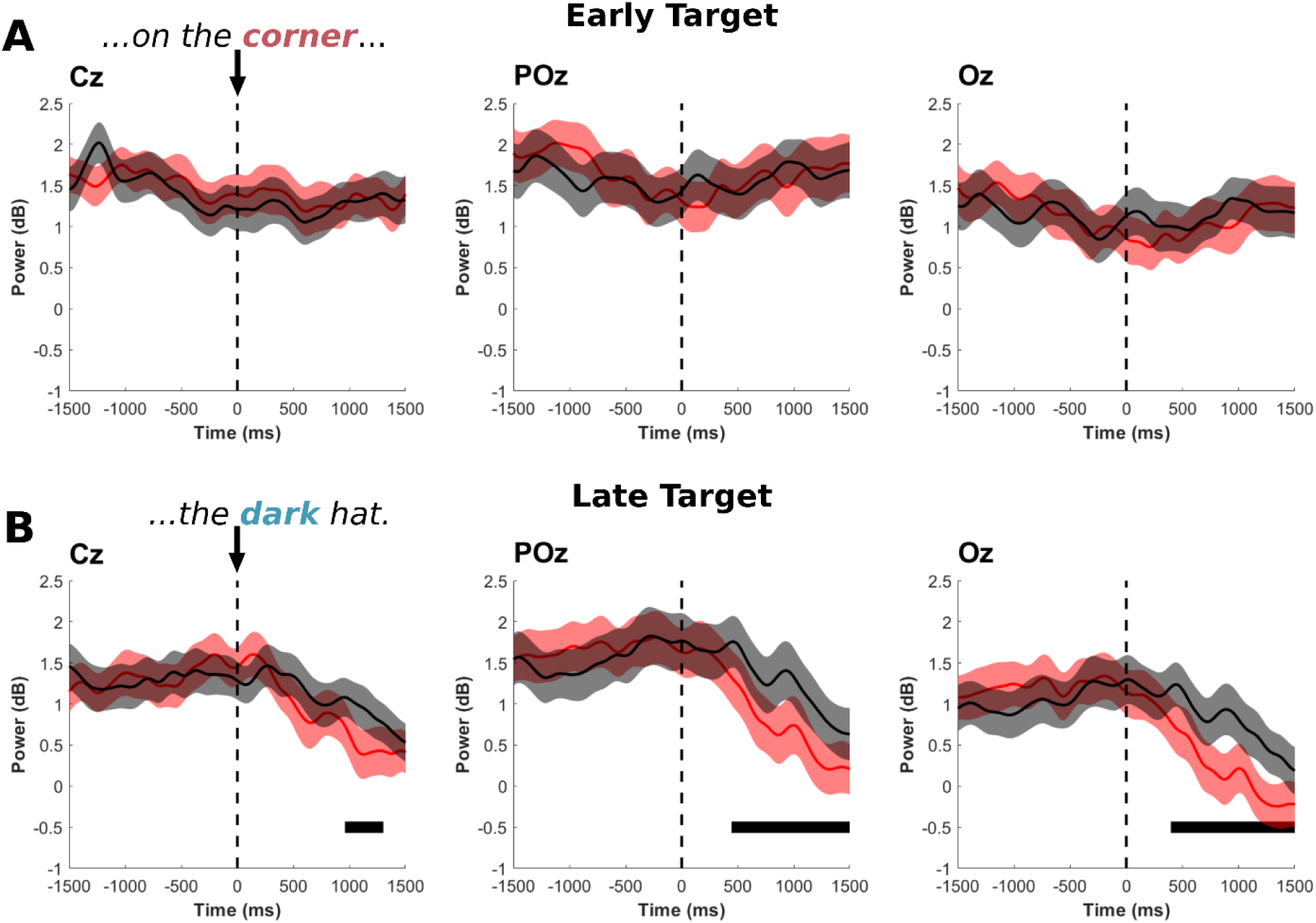
Power averaged over the alpha band (8-13 Hz), locked to the onset of the Early target word (A) and Late target word (B), for electrodes Cz, POz and Oz. Shaded areas represent Standard Error of the Mean. Horizontal black lines represent significant clusters.

Visualization of changes in spectral power relative to the 1000 ms baseline period prior to the question onset (Fig. 6) reveal that there was a marked increase in alpha power over parieto-occipital electrodes with the onset of the question (the first auditory stimulus of the trial), which then returned to baseline at the end of the sentence (the last auditory stimulus of the trial, on average ∼1140 ms after the Late target word onset). This fits with prior studies showing an increase in parieto-occipital alpha in preparation for an auditory stimulus (Fu et al., 2001), potentially reflecting a gating-by-inhibition mechanism whereby irrelevant visual information is suppressed in favor of auditory attention (Jensen & Mazaheri, 2010). Thus, we performed follow-up analyses on alpha power at midline parieto-occipital electrodes matching the observed scalp distribution of alpha power (CPz, Pz, POz, Oz).

**Figure 6.**
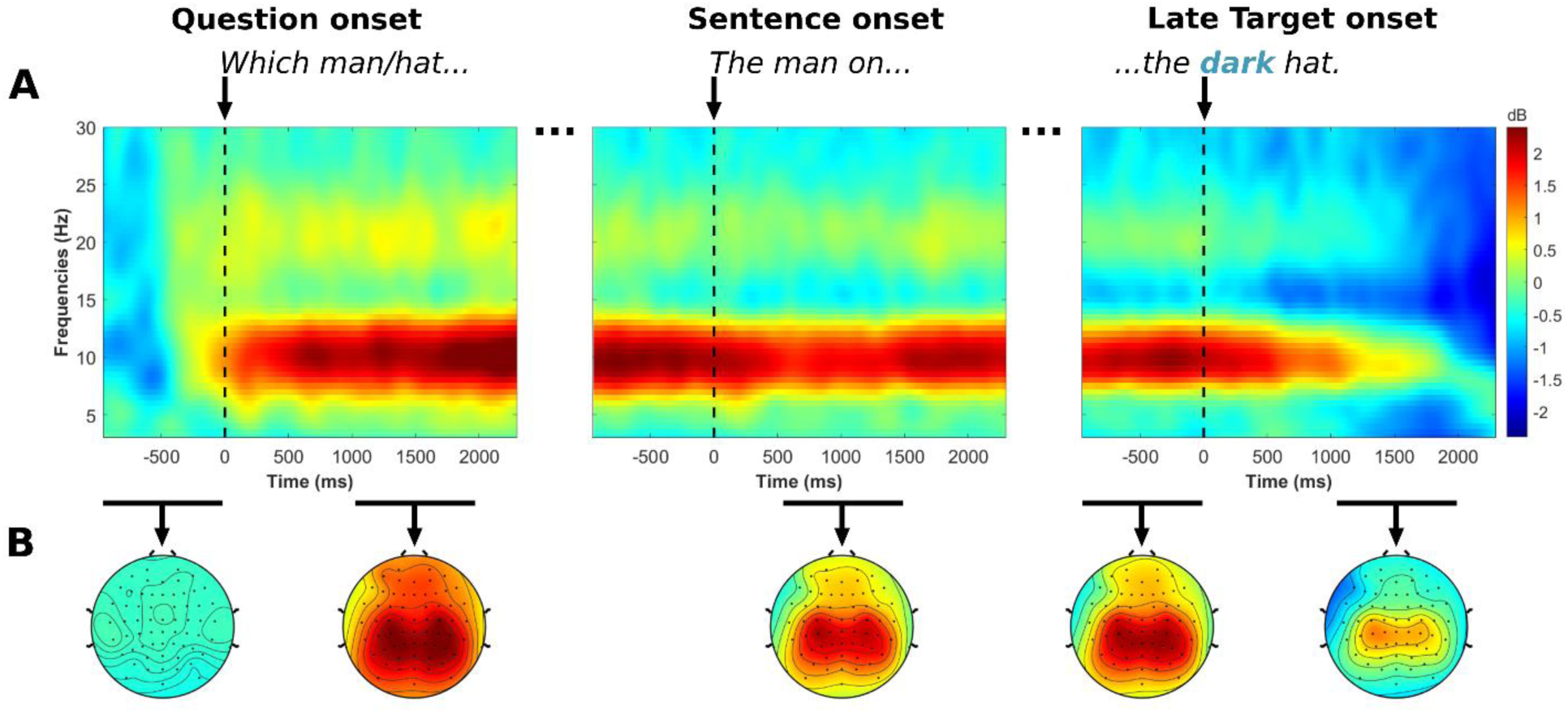
(A) Event-related spectral power at electrode POz throughout the trial, showing an increase in power over the alpha band (8-13 Hz) with the onset of the question and a return to baseline following the Late target word. (B) Scalp distribution of alpha power throughout the trial.

**Figure 6.**
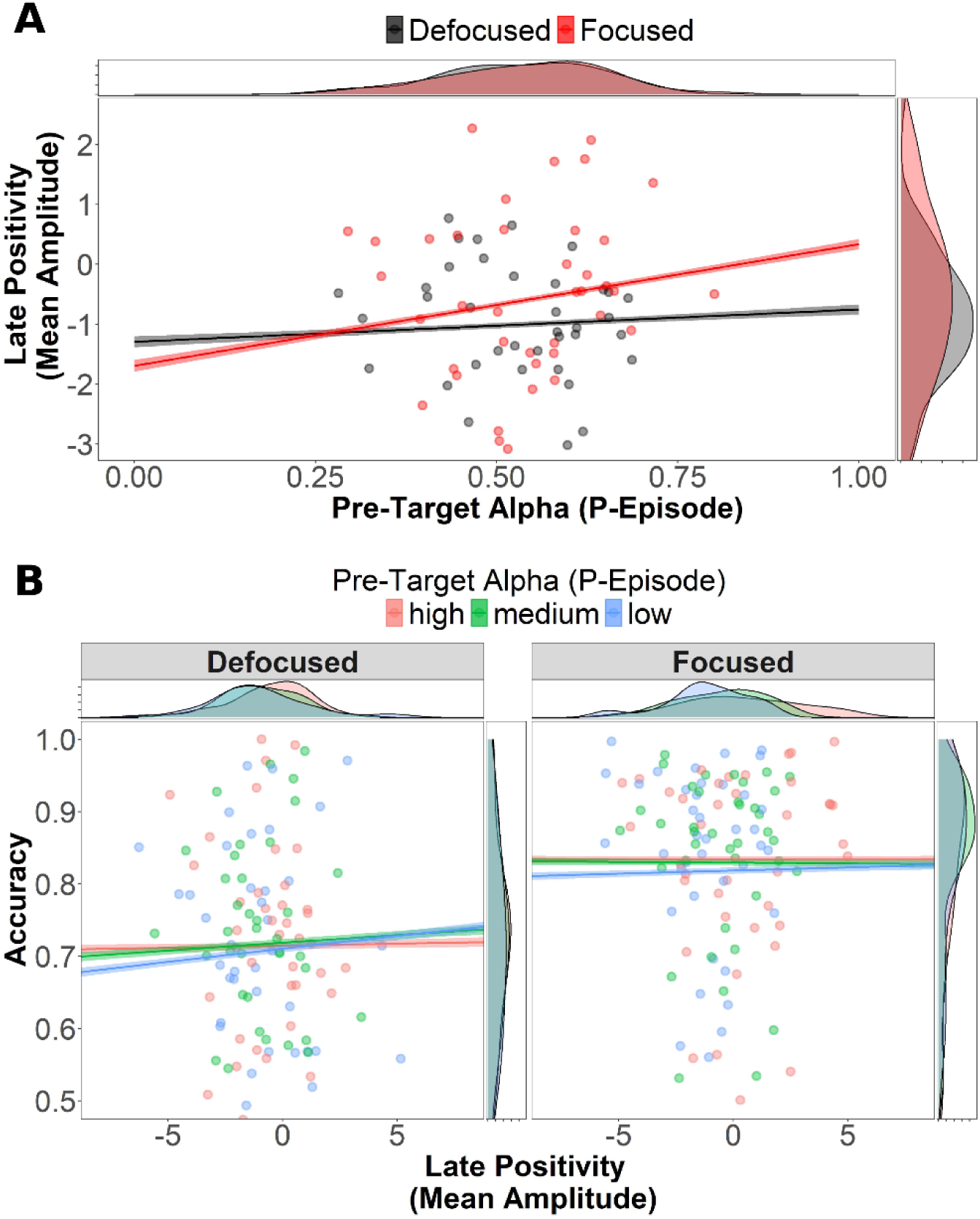
(A) Mean amplitude of the late positivity ERP (averaged over 600-1000 ms post-target) as a function of pre-target alpha (p-episode) and focus condition. Regression lines represent the relationship between mean amplitude and pre-target alpha, calculated from trial-level data. Individual data points and side density plots represent mean data for each participant, averaged across trials. (B) Memory test accuracy as a function of the mean amplitude of the late positivity ERP (averaged over 600-1000 ms post-target), pre-target alpha (p-episode) and focus condition. The y-axis is truncated at 0.5 for easier visualization. For visualization purposes only, pre-target alpha was turned to a categorical variable (high, medium, low; see legend), by splitting trials into three quantile-based groups – though analyses were conducted using a continuous measure of p-episode. Regression lines represent the relationship between mean amplitude and accuracy, calculated from trial-level data. Individual data points and side density plots represent mean data for each participant, averaged across trials. Greater accuracy was found for Focused words, and as a function of mean amplitude of the late positivity. The Dm effect of the late positivity on accuracy was stronger in the Defocused condition, and for trials with lower pre-target alpha (less attentional preparation).

While there was a trend for higher pre-target alpha (increased auditory attention through the inhibition of visual information) prior to Focused target words at electrodes POz and Oz, cluster-based permutations did not reveal any significant clusters prior to target onset. Instead, we found lower post-target alpha following the Focused Late target word, resulting in a significant cluster between 446 - 1500 ms post-target for electrode POz, and 397 - 1500 ms for Oz. This effect may reflect a faster return to baseline alpha power (releasing the occipital inhibition) after hearing the Focused Late target, in preparation for the behavioral task of answering the question out loud (which was indicated visually through a question mark on the screen). We return to this interpretation in the *Discussion*.

Given that any potential pre-target effects of focus on alpha may have been obscured by the jitter naturally present in our stimuli due sentence-level differences in duration and number of words, we also performed a follow-up analysis on pre-target alpha p-episode to overcome this issue (see *Method*). The Bayesian ordered-beta regression found several effects of electrode cluster, matching the observed scalp distribution of alpha power, such that p-episode was greater at Posterior clusters (see Table S4 for model output^4^). However, it did not reveal an effect of focus for either the Early target (*95% CI* = [-0.08, 0.06]) or the Late target (*95% CI* = [−0.01, 0.14]). To test the strength of the evidence for the null hypothesis, we calculated Bayes factors for all effects using the *bayestestR* package in R (Makowski et al., 2019). The Bayes factor for the effect of focus was 0.007 for the Early target and 0.026 for the Late target, indicating strong evidence in favor of the null hypothesis (Quintana & Williams, 2018). This confirms that this null effect was not due to lack of statistical power. Thus, based on both this analysis and the cluster-based permutations on alpha power, we conclude that focus condition did not have a significant effect on pre-target alpha in our study. Nonetheless, we tested whether pre-target alpha affected the ERP and behavioral responses on a trial-by-trial basis, under the assumption that – contrary to Boudewyn and Carter (2018) – *higher* pre-target alpha reflected *greater* auditory attention (through greater inhibition of visual information).

In summary, we did not find evidence that discourse focus modulated pre-target alpha power, either through cluster-based permutation testing or p-episode analysis, and Bayesian statistics provided strong support for the null hypothesis. While a post-target reduction in alpha power was observed following Focused Late targets over posterior electrodes, this effect may reflect preparatory processes for the upcoming behavioral response rather than anticipatory attention to focused words. Thus, contrary to our predictions and previous findings, focus did not significantly influence preparatory attention in this study.

### ERPs as a function of pre-target alpha

We tested whether the mean amplitude of the late positivity varied as a function of pre-target alpha on a trial-by-trial basis, using p-episode during the 1500 ms prior to each target word as our alpha measure. Data were aggregated across the Early and Late target words. There was a significant effect of alpha, such that greater pre-target alpha corresponded to a more positive late positivity (β = 1.35, *SE* = 0.06, *p* < 0.0001; Fig. 7), while the effect of focus was no longer significant (*p* > 0.05). There was also a significant interaction between pre-target alpha and focus, such that the effect of pre-target alpha on mean amplitude was larger for Focused words (β = −0.87, *SE* = 0.06, *p* < 0.0001), suggesting that trial-level variability in preparatory attention mediated the effect of focus on the late positivity.

### Behavior as a function of ERPs and pre-target alpha

The logistic mixed effects model on memory test accuracy, aggregated over the Early and Late targets, found that accuracy was higher for words that were Focused (β = −0.35, *SE* = 0.007, *p* < 0.0001), similarly to the prior behavioral analyses (where this effect was shown as a significant interaction between focus position and target position; see *Behavioral results*). Accuracy was also higher for words that elicited a more positive late positivity (mean amplitude over 600-1000 ms; β = 0.11, *SE* = 0.006, *p* < 0.0001; Fig. 6). Given the latency and scalp distribution of this ERP component, we interpret this as a “difference due to memory” (Dm) effect (Friedman & Johnson Jr., 2000; Mecklinger & Kamp, 2023; Paller et al., 1987). In particular, our effect matches prior reports of a centro-parietal positivity at a similar latency for words that were later remembered (Mecklinger & Kamp, 2023; Schott et al., 2002). The effect of mean amplitude on accuracy also interacted with focus, such that there was a larger effect for words that were Defocused (β = 0.06, *SE* = 0.006, *p* < 0.0001). This might be partly due to a ceiling effect in the Focused condition, but it also matches prior findings indicating that the Dm effect is larger for conditions of shallow processing (Schott et al., 2002).

There was no effect of pre-target alpha on accuracy (*p* > 0.05), but alpha significantly interacted with the effect of ERP mean amplitude (β = −0.06, *SE* = 0.006, *p* < 0.0001), such that mean amplitude had a larger effect on accuracy for trials with lower pre-target alpha. The three-way interaction between focus, mean amplitude and pre-target alpha was not significant. We checked for multicollinearity using the *performance* package in R (Lüdecke et al., 2021), which revealed that all effects had a variance inflation factor (VIF) below 5, indicating low multicollinearity.

Overall, this analysis revealed that trial-level differences in the amplitude of the late positivity predicted subsequent accuracy on the memory test (Dm effect), and that this effect was largest for words receiving shallow processing (when defocused by the preceding question) and less attentional preparation (lower pre-target alpha).

## Discussion

This study investigated the predominant hypothesis that listeners pre-allocate attention to the timing of focused, novel information. Attention has been invoked as the explanation for prior behavioral and ERP effects showing faster reaction times, deeper semantic and syntactic processing, and greater memory for words expected to be focused based on the discourse context, prosody and syntax (Akker & Cutler, 2003; Beier & Ferreira, 2022; Cutler & Fodor, 1979; Kristensen et al., 2013a; Wang et al., 2014; Yang et al., 2019). Here, we tested whether there is any evidence for attentional preparation prior to words focused or defocused by a preceding discourse question, measured through the dynamics of EEG alpha neural activity.

Alpha activity tracks preparatory attention in non-linguistic studies of temporal attention (Breska & Deouell, 2017b, 2017a; Nobre & Van Ede, 2018), and has been shown to reflect auditory attention to speech (Boudewyn et al., 2015, 2017; Boudewyn & Carter, 2018; Dimitrijevic et al., 2017; Wöstmann et al., 2016, 2017).

Contrary to our hypothesis based on this prior work, we did not find an effect of discourse focus on pre-target alpha power. We found that alpha power over posterior sites increased relative to baseline during the auditory presentation of the speech stimuli, returning to baseline after the stimulus ended. This pattern may reflect increased auditory attention through gating-by-inhibition, whereby irrelevant visual information processed in the occipital cortex is suppressed in favor of auditory processing (Jensen & Mazaheri, 2010). While we observed a pattern for greater pre-target alpha prior to focused words at posterior electrodes (which would suggest greater auditory attention through greater visual inhibition), this difference was not significant in our analyses. Instead, we found a faster return to baseline alpha at the end of the sentence when the late target word had been focused. This effect may reflect a release from occipital inhibition (and consequently an increase in visual attention) in preparation for the visual appearance of the question mark, which indicated whether participants would need to respond to the discourse question by verbally repeating the focused target word. After hearing the focused late target word (the answer to the question), participants prepared for this visual cue. Thus, while this was likely a task-specific effect, it shows that we could measure an effect of preparatory visual attention on alpha similar to those found in prior non-linguistic studies (Breska & Deouell, 2017b, 2017a); this further underscores that the null effect of focus on pre-target alpha was not due simply to lack of statistical power. However, while pre-target alpha did not vary as a function of discourse focus, we found that trial-level fluctuations in alpha power influenced the processing and memory for the target words, which we turn to next.

While discourse focus did not influence the amplitude of the N400, contrary to our predictions, focused words elicited a late centro-parietal positivity, and the amplitude of this positivity predicted whether the word would later be remembered. Based on this relationship, we interpret this as a “difference due to memory” (Dm) effect—an ERP signature associated with successful memory encoding (Friedman & Johnson Jr., 2000; Mecklinger & Kamp, 2023; Paller et al., 1987). Dm effects, also known as subsequent memory effects (SMEs), are differences in neural activity during encoding that depend on whether an item is later remembered or forgotten. Although their timing and scalp distribution can vary depending on task demands and retrieval format, our findings align with a well-documented positive-going parietal Dm effect, which has been linked to familiarity-based memory (i.e. as opposed to recollection; Meng et al., 2014; Yonelinas et al., 2010). This interpretation is consistent with our memory test design: participants selected the missing word from four options, a format that allows for recognition based on familiarity rather than having to freely recall the missing word.

This positivity is thought to reflect the formation of item-specific memory traces through the integration of new information with prior semantic knowledge (Mecklinger & Kamp, 2023). This interpretation fits our task structure, in which focused words were processed within a meaningful sentence context that encouraged semantic integration. Somewhat surprisingly, however, the Dm effect—defined as the ERP difference between remembered and forgotten items—was larger for defocused words. While defocused items are typically assumed to receive shallower processing (Wang et al., 2011), this finding is consistent with prior reports that Dm effects can be more pronounced under shallow encoding conditions (Schott et al., 2002). In such cases, memory success depends more on trial-by-trial variation in attention or engagement, leading to larger ERP differences between trials that do and do not result in successful encoding.

Supporting this view, we found that trial-level variability in pre-target alpha power influenced both the amplitude of the late positivity and its relationship to memory. Specifically, greater pre-target alpha—interpreted as greater preparatory auditory attention via inhibition of visual input—was associated with a larger late positivity, especially for focused words. This suggests that when a word was in focus, participants were more likely to engage attentional resources in anticipation of encoding, enhancing processing of the target word. While pre-target alpha did not predict memory accuracy directly, lower pre-target alpha was associated with a larger difference in the late positivity between remembered and forgotten items. This again matches prior findings that Dm effects tend to be stronger under shallow processing (Schott et al., 2002), where attentional variability plays a larger role in determining memory outcomes.

Together, these results suggest that both discourse focus and attentional state shape encoding success, but that the strength and variability of these effects differ depending on the depth of processing and the degree of preparatory attention.

### Role of alpha in preparatory attention to speech

While prior work from our lab found that increased alpha power during the discourse context predicted *reduced* sensitivity to referential ambiguity (as reflected in a smaller Nref; Boudewyn et al., 2015), the present findings show that greater alpha power prior to target words was associated with *enhanced* encoding and better subsequent memory. Although these findings may appear contradictory, they likely reflect different functional roles of alpha activity across task contexts. In the earlier study, increased alpha during the discourse context likely reflected reduced attention to the unfolding linguistic input, impairing the formation of distinct referential representations. In contrast, in the current study, increased alpha—particularly over posterior electrodes—preceded the target word itself, and likely reflected a gating mechanism, whereby visual input was inhibited to prioritize auditory processing (Fu et al., 2001; Jensen & Mazaheri, 2010). This interpretation is supported by the scalp distribution of the effect and by the preparatory nature of the pre-target window.

Together, these findings illustrate that alpha power is not a unitary marker of “attention” or “engagement,” but rather reflects flexible, context-sensitive neural filtering mechanisms.

Depending on task demands, increased alpha can either hinder processing (by reflecting disengagement from critical context) or facilitate it (by suppressing irrelevant input to optimize target encoding). This matches prior reports of both alpha synchronization and desynchronization elicited by attention during language comprehension (Dimitrijevic et al., 2017). Future work should further address how these different mechanisms vary as a function of the task and stimulus features.

Importantly, the fact that we did not see a difference in pre-target alpha as a function of discourse focus casts doubt on the common explanation for the effects of information structure on comprehension in terms of attention pre-allocation. In particular, our findings do not support the view that discourse questions guide attention to the timing to upcoming focused words through attentional mechanisms similar to those described in non-linguistic studies of temporal prediction and attention (Nobre & Van Ede, 2018). Instead, prior behavioral and ERP effects of focus could be explained by language-specific mechanisms, such as modulations in the strength and content of semantic predictions (Ferreira & Lowder, 2016). This does not necessarily deny the involvement of attention by information structure: rather than top-down attentional preparation, prior fMRI and ERP evidence for the engagement of attention may reflect a bottom-up response to salient information (e.g., elicited by pitch accents in Kristensen et al., 2013).

Nonetheless, discourse focus may have a more nuanced relationship with attentional preparation. Rather than varying as a function of focus, pre-target alpha appeared to reflect local trial-level variability in attention (similar to the attentional lapses in Boudewyn & Carter, 2018). Trial-by-trial differences in pre-target attention predicted post-target ERP responses associated with memory encoding, and this effect was strongest when words had been focused. Thus, discourse focus may determine the functional role of these trial-level fluctuations in attention, such that moments of greater preparatory attention are more likely to facilitate the encoding of focused information into memory.

### Limitations and future directions

A limitation of our design was that the exact timing of the early and late target words was necessarily jittered across trials, given the differing duration and number of words across sentences. This resembles naturalistic language processing, where the timing of a focused word may not be predictable to an exact degree. However, in this way our study departed from non-linguistic studies of temporal prediction, where a cue is typically associated with specific intervals of time. Thus, it is possible that the temporal predictions formed by the discourse questions in our study were not precise enough to be measurable through alpha power. We overcame this limitation through the use of the BOSC method to quantify pre-target alpha in terms of the percentage of timepoints containing above-threshold alpha regardless of its exact timing, which revealed trial-level effects of alpha on post-target responses. Nonetheless, future studies could employ a more controlled – albeit less natural – design where the timing of the target words is kept consistent across trials.

Another possibility is that the use of clear, read speech made comprehension too easy, such that effortful attentional pre-allocation was not required to complete the task. Future studies may test whether evidence for attentional preparation emerges when the intelligibility of speech is decreased, such as through a speech-in-noise task. Prior work shows that the comprehension of acoustically degraded speech leads to a decrease in alpha power corresponding to the amount of acoustic detail (Obleser & Weisz, 2012); combining this approach with our discourse focus manipulation will determine whether this effect is modulated by predictions about the timing of upcoming focused target words.

Further, while this study manipulated information structure through the discourse context, future studies should address whether preparatory attention is engaged by prosodic and syntactic cues to the timing of an upcoming focused word. While all of these cues were suggested to play a similar role by directing attention to focused words (Cutler & Fodor, 1979), it is possible that preparatory attentional effects are only elicited through prosody or syntactic focus. Additionally, studies have distinguished between ERP responses engaged by focus (i.e., through a discourse question) and by newness of information when the two are orthogonally manipulated (Chen et al., 2014). In our study, focused information was also always new to the listener; thus, dissociating between the two may lead to new insights into the role of focus on attentional preparation.

Finally, while we did not see an effect of focus on the N400 elicited by target words, our stimuli were not designed to lead to particularly predictable or unpredictable words. While prior work has shown that the N400 effect of semantic violations is enhanced for words focused through the discourse and prosody (Wang et al., 2011), future studies could test whether focus likewise affects the processing of words that were not predicted but that are plausible within the context, relative to predictable words. This is particularly relevant given that speakers are more likely to focus novel information (that is therefore less predictable, yet plausible). Further, as alternative explanations for the role of information structure on comprehension involve semantic prediction rather than attention allocation, a central question for future research is to determine how exactly the expectation of focus based on prior discourse, prosody and syntax interacts with semantic processing.

## Conclusions

While prior accounts have proposed that comprehenders strategically allocate attention to the anticipated timing of focused, informative input, our results suggest that such preparatory mechanisms may not consistently manifest in neural markers like alpha power—at least under the listening conditions examined here. This opens the possibility that the cognitive and neural effects of information structure may arise not from controlled pre-allocation of attention, but instead from language-specific processes (e.g., shifts in semantic prediction) or bottom-up attentional capture once the relevant input is encountered.

These findings raise important questions about how listeners prepare for and respond to focused information, and what mechanisms support the processing advantages typically associated with focus. Future work should examine these mechanisms under varied listening conditions (e.g., in the presence of background noise) and across different cues to information structure, such as prosody and syntactic construction, to determine when and how attention is modulated during comprehension.

## Acknowledgments

This work was supported by a Ruth L. Kirschstein Postdoctoral Individual National Research Service Award (F32) from the Eunice Kennedy Shriver National Institute of Child Health and Human Development awarded to EB (grant number: F32HD108937), as well as a Short-Term Research Grant from the German Academic Exchange Service (DAAD) awarded to EB. We would like to thank all members of the Mangun, Swaab, Miller, Breska and Oganian labs for their exceptional support and feedback, with particular thanks to Timothy Trammel for his help recording the stimuli and Daniel Comstock for his support with the analyses. We would also like to thank Fernanda Ferreira for her feedback at the early conceptual stages of this project, and Steve Luck for his useful comments. Lastly, we extend special gratitude to the hard work of the undergraduate research assistants who were involved in stimulus preparation and data collection, in alphabetical order: Grace Bell, Saya Davani, Sonakshi Khanna, Sophia Kinnear, Lindsay Li and Hao Ngo.

## Footnotes

1 Due to a technical issue with the apparatus, EOG data was not recorded for 13 out of 40 participants. No significant difference was found between participants with and without EOG data in any of the behavioral or EEG measures of interest.

2 We constrained this analysis to midline electrodes given that we did not have any a-priori hypotheses about lateralized effects.

3 For all models, in case of convergence issues or singular fit, the random effects structure was simplified by iteratively removing the random effect explaining the least amount of variance until the model converged (Barr et al., 2013).

4 For Bayesian models, we considered an effect to be “significant” when the 95% Credible Interval for the effect did not overlap with 0 (Nicenboim & Vasishth, 2016).

