## Supplementary Tables for "Discourse Focus and Memory Encoding: The Role of Trial-Level Alpha Power"

**Table S1**

*Mixed effects model output. Mean ERP amplitude at 300-500 ms for the filler items, as a function of semantic constraint and electrode cluster.*

| **Predictor** | **β** | ***SE*** | ***p*** | |
|---|---|---|---|---|
| (Intercept) | 0.07 | 0.29 | .805 | |
| Semantic Constraint (Low Constraint) | -2.26 | 0.40 | < .001 | *** |
| Cluster (Left Frontal) | -0.86 | 0.12 | < .001 | *** |
| Cluster (Left Posterior) | 0.48 | 0.13 | < .001 | *** |
| Cluster (Left Temporal) | -0.14 | 0.13 | .298 | |
| Cluster (Mid Frontal) | -0.59 | 0.13 | < .001 | *** |
| Cluster (Mid Posterior) | 0.47 | 0.13 | < .001 | *** |
| Cluster (Right Frontal) | -0.20 | 0.12 | .114 | |
| Cluster (Right Posterior) | 0.56 | 0.13 | < .001 | *** |
| Cluster (Right Temporal) | 0.14 | 0.13 | .302 | |
| Low Constraint × Left Frontal | 1.25 | 0.18 | < .001 | *** |
| Low Constraint × Left Posterior | 0.92 | 0.18 | < .001 | *** |
| Low Constraint × Left Temporal | 1.00 | 0.19 | < .001 | *** |
| Low Constraint × Mid Frontal | 0.99 | 0.19 | < .001 | *** |
| Low Constraint × Mid Posterior | 0.43 | 0.19 | .024 | * |
| Low Constraint × Right Frontal | 1.05 | 0.18 | < .001 | *** |
| Low Constraint × Right Posterior | 0.78 | 0.18 | < .001 | *** |
| Low Constraint × Right Temporal | 1.05 | 0.19 | < .001 | *** |

**Note.** Significance codes: *** $p < .001$, ** $p < .01$, * $p < .05$, . $p < .10$. Fixed effects were dummy-coded, using the High Constraint condition and the Central cluster as reference levels. *SE* = Standard Error.

**Table S2.**

*Mixed effects model output. Mean ERP amplitude at 300-500 ms for the Early and Late targets, as a function of focus and electrode cluster.*

| | Early Target | | | | Late Target | | | |
|---|---|---|---|---|---|---|---|---|
| **Predictor** | **β** | ***SE*** | ***p*** | | **β** | ***SE*** | ***p*** | |
| (Intercept) | -3.23 | 0.35 | < .001 | *** | -3.15 | 0.37 | < .001 | *** |
| Focus (Focused) | -0.36 | 0.32 | .258 | | -0.31 | 0.32 | .336 | |
| Cluster (Left Frontal) | 1.03 | 0.29 | < .001 | *** | 2.32 | 0.33 | < .001 | *** |
| Cluster (Left Posterior) | 2.18 | 0.30 | < .001 | *** | 1.09 | 0.34 | .002 | ** |
| Cluster (Left Temporal) | 1.45 | 0.31 | < .001 | *** | 1.37 | 0.35 | < .001 | *** |
| Cluster (Mid Frontal) | 0.98 | 0.31 | .003 | ** | 1.77 | 0.35 | < .001 | *** |
| Cluster (Mid Posterior) | 1.42 | 0.31 | < .001 | *** | 0.30 | 0.35 | .396 | |
| Cluster (Right Frontal) | 1.20 | 0.29 | < .001 | *** | 2.02 | 0.33 | < .001 | *** |
| Cluster (Right Posterior) | 2.00 | 0.30 | < .001 | *** | 0.86 | 0.34 | .014 | * |
| Cluster (Right Temporal) | 1.22 | 0.31 | < .001 | *** | 1.02 | 0.35 | .006 | ** |
| Focus × Left Frontal | -0.16 | 0.15 | .291 | | -0.96 | 0.16 | < .001 | *** |
| Focus × Left Posterior | 0.44 | 0.16 | .006 | ** | 0.90 | 0.16 | < .001 | *** |
| Focus × Left Temporal | 0.25 | 0.16 | .123 | | 0.07 | 0.17 | .688 | |
| Focus × Mid Frontal | -0.26 | 0.16 | .118 | | -0.71 | 0.17 | < .001 | *** |
| Focus × Mid Posterior | 0.14 | 0.16 | .379 | | 0.83 | 0.17 | < .001 | *** |
| Focus × Right Frontal | -0.01 | 0.15 | .965 | | -0.28 | 0.16 | .070 | . |
| Focus × Right Posterior | 0.09 | 0.16 | .563 | | 0.83 | 0.16 | < .001 | *** |
| Focus × Right Temporal | 0.03 | 0.16 | .853 | | 0.29 | 0.17 | .082 | . |

**Note.** Significance codes: *** $p < .001$, ** $p < .01$, * $p < .05$, . $p < .10$. Fixed effects were dummy-coded, using the Defocused condition and the Central cluster as reference levels. *SE* = Standard Error.

**Table S3.**

*Mixed effects model output. Mean ERP amplitude at 600-1000 ms for the Early and Late targets, as a function of focus and electrode cluster.*

| | Early Target | | | | Late Target | | | |
|---|---|---|---|---|---|---|---|---|
| **Predictor** | **β** | ***SE*** | ***p*** | | **β** | ***SE*** | ***p*** | |
| (Intercept) | -2.65 | 0.34 | < .001 | *** | -1.35 | 0.33 | < .001 | *** |
| Focus (Focused) | 0.70 | 0.39 | .079 | . | 1.08 | 0.40 | .009 | ** |
| Cluster (Left Frontal) | 0.78 | 0.12 | < .001 | *** | 0.35 | 0.13 | .006 | ** |
| Cluster (Left Posterior) | 2.52 | 0.13 | < .001 | *** | 1.46 | 0.13 | < .001 | *** |
| Cluster (Left Temporal) | 1.64 | 0.14 | < .001 | *** | 0.84 | 0.14 | < .001 | *** |
| Cluster (Mid Frontal) | 0.62 | 0.16 | < .001 | *** | 0.43 | 0.16 | .007 | ** |
| Cluster (Mid Posterior) | 1.63 | 0.14 | < .001 | *** | 1.07 | 0.14 | < .001 | *** |
| Cluster (Right Frontal) | 1.04 | 0.12 | < .001 | *** | 0.02 | 0.13 | .878 | |
| Cluster (Right Posterior) | 2.15 | 0.13 | < .001 | *** | 1.25 | 0.13 | < .001 | *** |
| Cluster (Right Temporal) | 1.37 | 0.14 | < .001 | *** | 0.45 | 0.14 | .001 | ** |
| Focus × Left Frontal | -1.10 | 0.18 | < .001 | *** | -2.13 | 0.18 | < .001 | *** |
| Focus × Left Posterior | -0.32 | 0.19 | .083 | . | 0.26 | 0.19 | .176 | |
| Focus × Left Temporal | -0.55 | 0.20 | .005 | ** | -0.62 | 0.20 | .002 | ** |
| Focus × Mid Frontal | -0.77 | 0.22 | < .001 | *** | -1.40 | 0.22 | < .001 | *** |
| Focus × Mid Posterior | -0.02 | 0.20 | .907 | | 0.49 | 0.20 | .013 | * |
| Focus × Right Frontal | -0.80 | 0.18 | < .001 | *** | -0.76 | 0.18 | < .001 | *** |
| Focus × Right Posterior | -0.18 | 0.19 | .348 | | 0.32 | 0.19 | .094 | . |
| Focus × Right Temporal | -0.31 | 0.20 | .112 | | 0.17 | 0.20 | .384 | |

**Note.** Significance codes: *** $p < .001$, ** $p < .01$, * $p < .05$, . $p < .10$. Fixed effects were dummy-coded, using the Defocused condition and the Central cluster as reference levels. *SE* = Standard Error.

**Table S4.**

*Bayesian ordered-beta mixed effects model output. Alpha p-episode at -1500-0 ms for the Early and Late targets, as a function of focus and electrode cluster.*

| | Early Target | | | Late Target | | |
|---|---|---|---|---|---|---|
| **Predictor** | **Estimate** | **Est. Error** | **95% CI** | **Estimate** | **Est. Error** | **95% CI** |
| Intercept | 0.22 | 0.06 | **[0.11 0.33]** | 0.17 | 0.06 | **[0.05 0.30]** |
| Focus (Focused) | -0.01 | 0.03 | [-0.08 0.06] | 0.06 | 0.04 | [-0.01 0.14] |
| Cluster (Left Frontal) | -0.22 | 0.01 | **[-0.25 -0.20]** | -0.25 | 0.01 | **[-0.28 -0.23]** |
| Cluster (Left Posterior) | 0.26 | 0.01 | **[0.24 0.29]** | 0.31 | 0.01 | **[0.28 0.33]** |
| Cluster (Left Temporal) | 0.05 | 0.01 | **[0.03 0.08]** | 0.05 | 0.01 | **[0.02 0.08]** |
| Cluster (Mid Frontal) | -0.29 | 0.01 | **[-0.31 -0.26]** | -0.30 | 0.01 | **[-0.33 -0.27]** |
| Cluster (Mid Posterior) | 0.26 | 0.01 | **[0.23 0.29]** | 0.33 | 0.01 | **[0.30 0.36]** |
| Cluster (Right Frontal) | -0.28 | 0.01 | **[-0.30 -0.25]** | -0.28 | 0.01 | **[-0.30 -0.25]** |
| Cluster (Right Posterior) | 0.26 | 0.01 | **[0.23 0.29]** | 0.31 | 0.01 | **[0.28 0.33]** |
| Cluster (Right Temporal) | 0.01 | 0.01 | [-0.01 0.04] | 0.02 | 0.01 | [-0.01 0.05] |
| Focus × Left Frontal | 0.05 | 0.02 | **[0.01 0.08]** | -0.00 | 0.02 | [-0.04 0.03] |
| Focus × Left Posterior | 0.03 | 0.02 | [-0.01 0.06] | -0.02 | 0.02 | [-0.06 0.01] |
| Focus × Left Temporal | 0.01 | 0.02 | [-0.02 0.05] | -0.02 | 0.02 | [-0.06 0.01] |
| Focus × Mid Frontal | 0.04 | 0.02 | **[0.00 0.08]** | -0.00 | 0.02 | [-0.04 0.04] |
| Focus × Mid Posterior | 0.03 | 0.02 | [-0.01 0.07] | -0.01 | 0.02 | [-0.05 0.03] |
| Focus × Right Frontal | 0.05 | 0.02 | **[0.02 0.09]** | -0.02 | 0.02 | [-0.06 0.01] |
| Focus × Right Posterior | 0.03 | 0.02 | [-0.01 0.06] | 0.00 | 0.02 | [-0.03 0.04] |
| Focus × Right Temporal | 0.04 | 0.02 | **[0.00 0.08]** | -0.02 | 0.02 | [-0.06 0.02] |

**Note.** 95% Credible Intervals that do not overlap with 0 are bolded. Fixed effects were dummy-coded, using the Defocused condition and the Central cluster as reference levels. CI = Credible Interval.
